# Reconstruction of Three-Dimensional Trajectories of Honeybees Flying in High-Density Aerial Environments

**DOI:** 10.1101/2022.11.24.517808

**Authors:** Mandiyam Yadav Mahadeeswara, Mandyam V Srinivasan

## Abstract

So far, relatively few studies have investigated the flight of insects moving in experimental conditions approximating natural scenes. This is mainly because of the technical difficulties involved in detecting and tracking individual insects in three dimensions in outdoor scenarios, as well as matching views of corresponding insects in stereo images for trajectory reconstruction (the so-called ‘Correspondence Problem’). In this study, we describe the methods we have developed to track and unambiguously reconstruct the trajectories and body orientations of a large number of bees flying in close proximity in a ‘bee cloud’, using just two high-speed video cameras configured as a stereo pair in a semi-outdoor setting. Using these methods, two separate bee clouds were filmed and the data were analysed to reconstruct the three-dimensional trajectories, including the head and tail positions, of a total of 382 bees. This dataset should enable future analysis of the movement characteristics of bees flying in a dense environment, as well as uncover potential strategies for mid-air collision avoidance.

## BACKGROUND AND SUMMARY

Relatively little is known about the flight characteristics and strategies employed by freely flying insects in an outdoor environment which, in general, is highly dynamic. Studies, conducted in indoor laboratory environments, have investigated the flight of honeybees and bumblebees through narrow corridors^1–5^ (see review^6^), gaps or apertures^7–10^, and collision avoidance and landing behaviour of fruitflies^11,12^. Yet, only a handful of studies have examined the behaviour of honeybees flying freely in a dynamically changing setting^13–15^ that approximates the more realistic scenario encountered by a foraging bee. Two primary reasons for this are: (a) the challenges of recording data in natural, uncontrolled scenes^16^ and (b) the difficulties associated with customizing the available tracking programs to fit one’s study requirements.

Algorithms have been developed for tracking multiple flying insects (mosquitoes^17^, midges^18–20^, honeybees^21,22^, fruitflies^23,24^ and birds^25–27^). Researchers have also developed tracking algorithms to reconstruct trajectories of freely flying *Drosophila* in virtual reality (VR) based systems^28–30^. In a recent study, Pannequin *et al* ^31^ used a SpiderCam^32^ setup, referred to as a ‘lab on cables’, to track the flight of a single freely flying moth. Most techniques track the approximate 3D position of the centre of gravity (CG) of the whole insect from each stereo pair by computing the CG of the pixels representing the insect in each camera image^20,33–35^. In practice, however, it is a challenge to track and accurately reconstruct the head and tail positions accurately and unambiguously from each pair of video frames, without any markers on the insect, especially when multiple insects are being filmed. Most of the studies conducted so far suffer from both of the following shortcomings: (a) usage of bulky and costly experimental apparatus, setups and methods which cannot be extended to tracking multiple flying insects in an outdoor setting^23,28,30,31,36–38^, and (b) the final digitised data often does not provide information on body orientation of the insects (or birds) flying in the group^17,18,21,22,29^.

As a step towards examining complex flight behaviours in a randomly changing quasi-natural setting, we have created a test scenario in a semi-outdoor environment, consisting of a cloud of loitering honeybees flying past one another with the goal of entering their hive, whose entrance has been temporarily blocked. We term this test scenario a ‘bee cloud’. Bees then fly in the vicinity of the hive entrance, and this scenario offers a valuable opportunity to study the behaviour of bees flying in a dense, dynamic aerial environment which could also approximate other dynamic scenarios such as dandelion seeds moving in the air.

The study also develops a tracking program that reliably detects, tracks and reconstructs the three-dimensional trajectories and the body orientations of a number of bees flying in a ‘bee cloud’. The program is capable of successfully dealing with some of the challenges, such as occlusions, that are encountered in a multi-object tracking scenario. Furthermore, the problem of correspondence matching is solved in a simple way by using just two cameras. The accuracies of two resulting stereomatching algorithms are assessed, compared and validated using a ‘ground truth’ control experiment.

In summary, we present a dataset that provides a full characterization of two bee cloud events captured in a semi outdoor environment, which can be used to visualize and study the movements and flight manoeuvres of a number of bees flying in close proximity. To our knowledge, our study is the first of its kind to (a) provide a dataset on bee clouds and (b) develop a reliable, error-free stereo-matching procedure to derive accurate three-dimensional trajectories of all of the bees flying in the cloud with high spatial and temporal resolution. This dataset should enable analysis of the movement characteristics of bees flying in a dense environment, and, potentially, strategies for mid-air collision avoidance.

## METHODS

### 1) EXPERIMENTAL ARENA

An independent honeybee (*Apis mellifera*) colony was stationed and maintained on the rooftop of our institute’s building. The bee colony was non-captive in nature, which provided unconditional access to the bees to forage in the surrounding environment. An image of the beehive on the rooftop of the building is shown in Fig. 1a. Three days before data collection, three rectangular white corflute sheets, of dimension 1.96 m (w) × 6 m (h), were erected on the three sides of the beehive (to its left, right & behind the hive entrance). The floor was covered with a sheet of white corflute. This created an arena in the form of a rectangular cuboid with white internal faces, against which the bees could be filmed in high contrast. A schematic illustration of the structure of the arena and the stereo video cameras is given in Fig. 1b.

**Figure 1.**
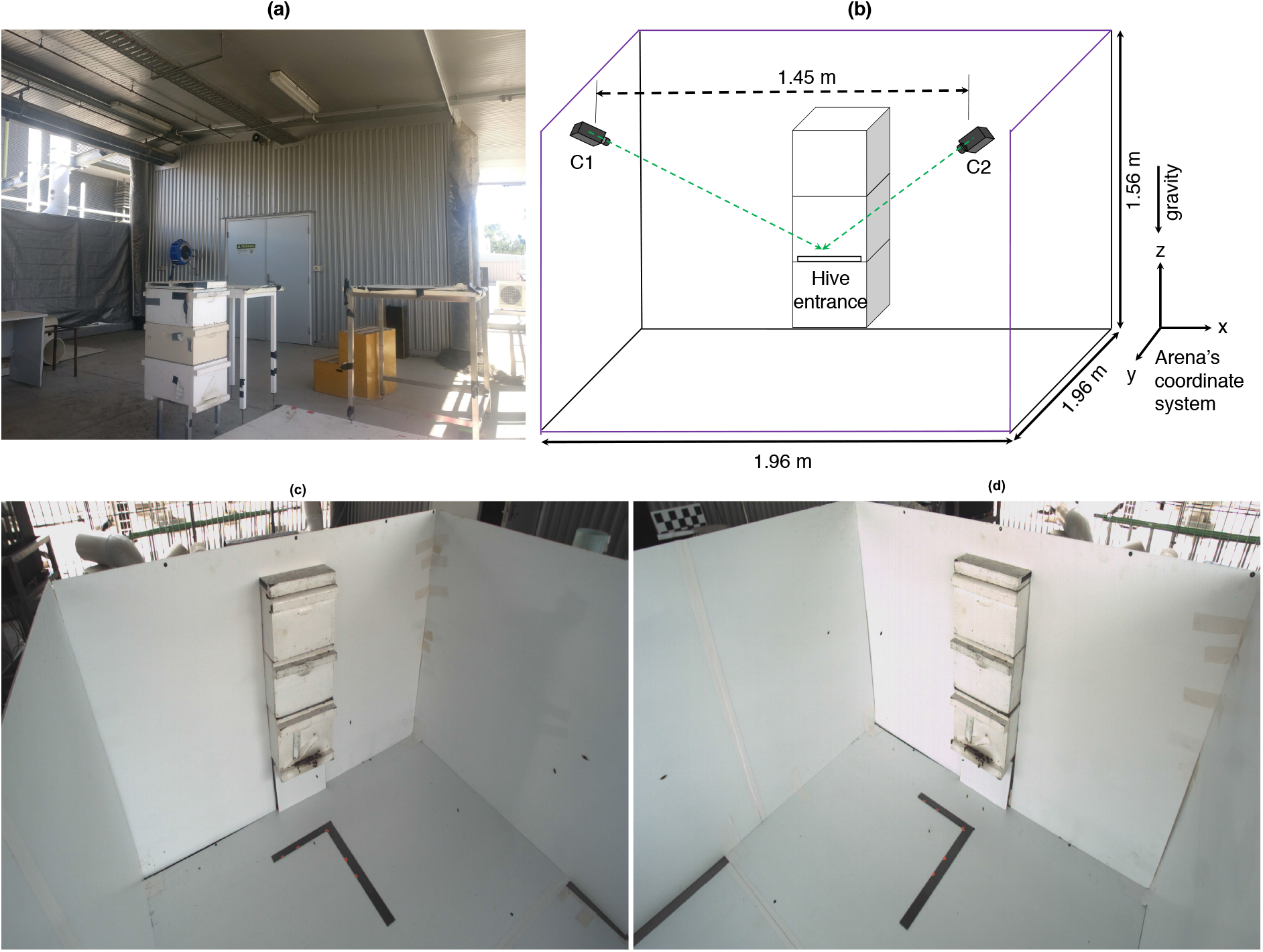
(a) Image of the beehive on the building’s rooftop. (b) Schematic diagram illustrating the structure of the experimental arena and the arrangement of the video cameras used in the study. The dashed green lines indicate the optical axis of each camera. The boundaries of the arena are shown in purple. (c,d) Stereo images of the arena, captured by the two cameras before commencing the experiment. The L-shaped square ruler was used to provide a reference x, y axis for calibration purposes.

After setting up the arena walls, we allowed the bees to get accustomed to the new arena for the next 3 days. Experiments were commenced from the 4^th^ day onwards. Bees flying in the arena were filmed using two synchronized, high-speed cameras (Emergent Vision Technologies), positioned as shown in Fig. 1b. Views of the scene from the two cameras are given in Fig. 1c and Fig. 1d.

Video recordings were made at a frame rate of 200 fps, at a pixel resolution of 2048×1024. The two cameras were separated by a distance of 1.45m horizontally and the cameras were mounted on tripods, at a height of approximately 1.5m above the ground. Each camera was equipped with a 6mm fixed focal length lens (VS-0618H1).

We calibrated the camera configuration using the Jean-Yves Bouguet camera calibration toolbox^39^ to obtain the intrinsic and extrinsic parameters of the two cameras. After these parameters were obtained, we set up a bee cloud.

#### GENERATING A BEE CLOUD

We temporarily blocked the hive entrance with a piece of white rectangular corflute. Because of this, the returning foraging bees were unable to enter the hive and loitered in front of the hive entrance forming a cloud shaped aggregation, which we term a ‘bee cloud’. Camera recordings were commenced about 30 seconds after blocking the hive entrance, by which time a sufficiently large number of foragers had returned to the area near the hive entrance to form a dense bee cloud. Each filming session lasted for a maximum of about 10 seconds, after which the hive entrance was reopened to allow the loitering bees to re-enter their hive.

The video recordings were usually made around 12 noon, when the foraging activity was relatively high.

The two datasets presented in this study are from recordings of 7.9 and 7.0 seconds, respectively. The two high resolution bee cloud videos can be accessed from Data Citation 1.

A few points to note are:

1. The hive entrance was not closed for more than 3 minutes continuously, as prolonged closure could disturb the bees’ foraging activity and induce them to swarm.
2. After recording a bee cloud event, an interval of at least 30 minutes was allowed before creating a subsequent bee cloud. This interval enabled the returning foragers to re-enter the hive, deposit their nectar/pollen, and resume their normal foraging activity.

#### ANALYSIS OF VIDEO DATA

Analysis of the video data commenced with the detection and tracking of the head and tail positions of bees, frame by frame, in the video sequences captured by the two cameras. Manual digitisation of video data with high frame rates and high spatial resolution is a tedious and time-consuming process. Therefore, we developed a semi-automatic tracking program for digitising the head and tail locations of individual bee images in the two camera views. The detection and tracking algorithm, and the methods for dealing with various complicating contingencies, are described below.

### 2) DETECTION AND TRACKING

The detection and tracking algorithm basically performs the following 5 steps:

1. The raw footage is initially processed to obtain an image of the background of the scene by averaging all the frames in the video. The averaging can be performed using ‘ImageJ’ (ImageJ - https://imagej.nih.gov/ij/) software, or by a custom-designed Matlab (MathWorks-https://au.mathworks.com/products/matlab.html) program. This results in an image, for each camera view, that contains only the background, without any bees.
2. As a second step, ‘image differencing’ is performed between each video frame and the background image, to detect all of the bees in each frame of the video sequence. This is done separately for the two video sequences. The image of each bee then appears as a dark blob against a grey, featureless background. The video frames are then binarized to set all pixels within each bee image to black, and all background pixels to white. Next, the boundary of the image of each bee is computed using the ‘bwboundaries’ command in Matlab. This command returns the coordinates of all of the pixels that comprise the contour of the image of each bee.
3. The boundary contour information is fitted to an ellipse using a customized open source program^40^ which performs a least-squares fit. This program returns the basic properties of the fitted ellipse - such as the centre of the ellipse, and the length and orientation of the major axis (See Fig. 2). These parameters are used to determine the pixel coordinates of the centroid and the head and tail of the bee’s image (The procedure for identifying the head and the tail is detailed below).
4. For tracking the image (head and tail coordinates) of a particular bee through the video, the user only has click on the bee image in the first frame. The developed program then delivers the head and tail coordinates of the selected bee using the logic explained below. Consider a bee that is moving in the direction indicated by the green arrow in Fig. 2. From the images of the bee in two consecutive frames (frame 1 and frame 2, shown as fitted ellipses), one can readily infer, through visual inspection, that the head and tail positions of the bee in the two frames are (*H*_1_, *T*_1_) and (*H*_2_, *T*_2_) respectively, assuming that the bee is moving forwards. The logic for automating this identification in software is as follows. If the bee is moving forward with its body pointed close to the direction of flight, then the distance between the tail of the bee in frame 1 (*T*_1_) and the head of the same bee in frame 2 (*H*_2_) will be larger than the distances between all of the head-tail combinations: that is, 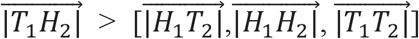 (see Fig. 2). This enables determination of the approximate flight direction as the direction of the vector 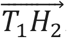. Given this flight direction, the head and tail locations in frame 1 are identified as follows. We compute the angles *θ*_1_ and *θ*_2_ between the flight direction vector 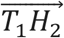 and the two possible orientations of the body axis, represented by the vectors 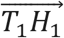 and 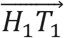, respectively (note that *θ*_1_ + *θ*_2_ = 180°). The vector associated with the smaller angle is taken to indicate the identities of the head and tail, as the body is more likely to be pointing toward the direction of movement, rather than in the opposite direction. Accordingly, in the example of Fig. 2, *H*_1_ and *T*_1_ are assigned as indicated by 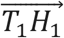, because *θ*_1_ < *θ*_2_. Once *H*_1_ and *T*_1_ are identified, *H*_2_ and *T*_2_ in frame 2 are identified by selecting the head-tail configuration that is associated with the smaller change in body orientation from frame 1 to frame 2: It is very unlikely that the body rotates by more than 90° in one interframe interval, given the high frame rate of 200/s. The validity of this assumption and the accuracy of this procedure are confirmed in the results presented later below. The areas of the ellipses fitted to the bee in the first two frames are computed and stored for subsequent use, for the purpose of detecting and resolving occlusions. This is described later below. When a bee moves backwards in the first two frames, the estimated head and tail positions obtained using the principle explained above would be reversed. When such a scenario arises, we simply swap the head-tail positions in the second frame, and run the tracking program as before to complete the tracking of that bee.
5. To successfully track a selected bee or a batch of bees from the third frame onwards, we initially predict the head and tail positions of the selected bee in the third frame by performing a simple 2-point linear extrapolation, based on the head and tail positions of the selected bee in the first two frames. We then search in the third frame for any bees whose centre of gravity (mid-point of the fitted ellipse) is within a radius of 20 pixels from the extrapolated head position of the selected bee. If more than one bee image is found in the search region, the orientation of the bee image tracked in frames 1 and 2 is used to determine the matching bee in frame 3. This selection is based on the reasoning that, given the relatively high frame rate of 200/s, bees are not likely to change their body orientation substantially between frames. The bee image chosen in frame 3 is therefore the one that is associated with the smallest change in the body orientation (orientation of the head-tail axis), and this is used to identify the positions of the head and tail of the bee’s image in frame 3. This two-layered process of (a) extrapolation and (b)matching of body orientation is repeated for each new frame, to track the image of each bee sequentially through all the video frames in which it is visible. The head and tail co-ordinates of each bee are also tagged with the frame number. As we shall describe later, this enables correspondence matching of the images of each bee in the image sequences captured by the two video cameras.

**Figure 2.**
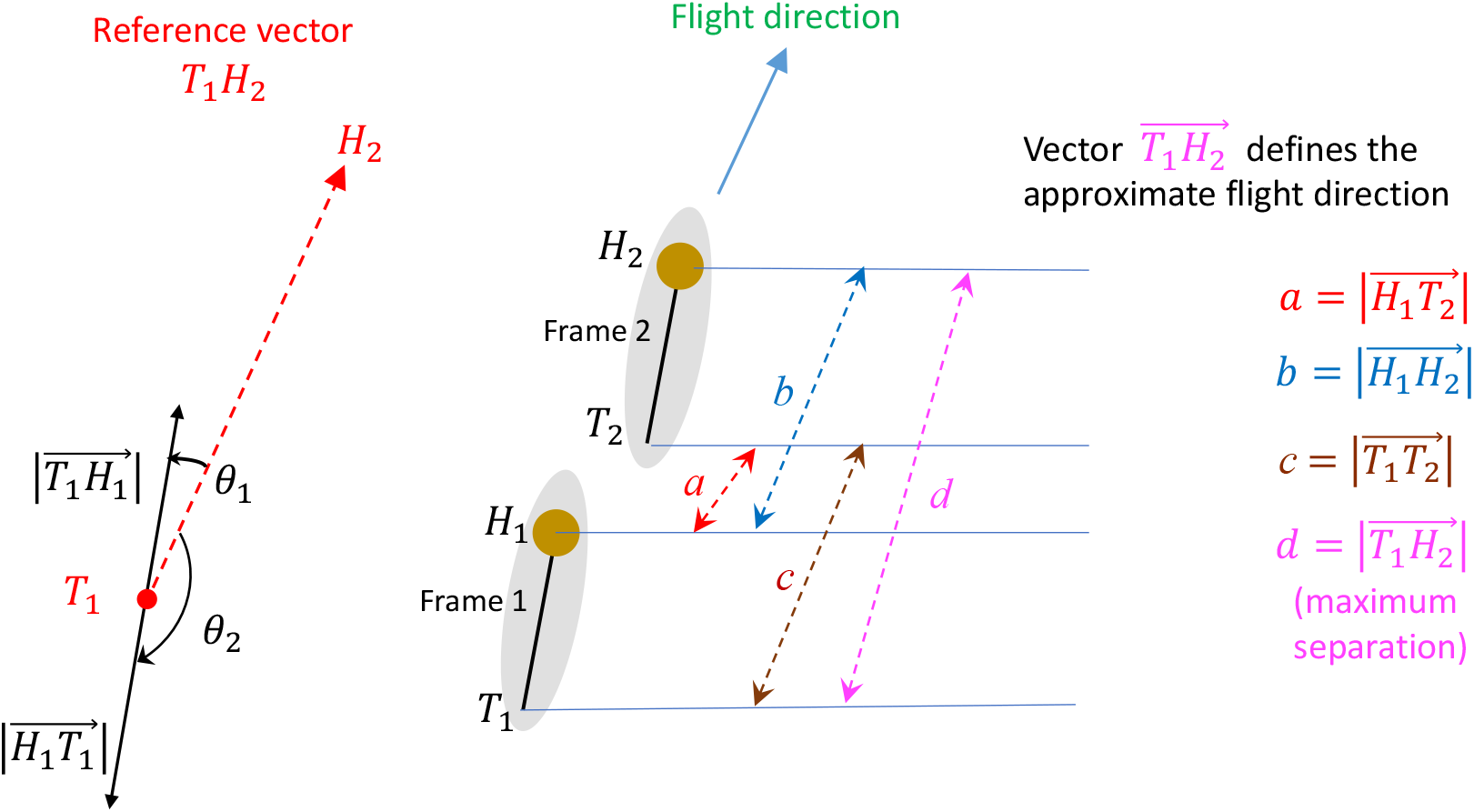
Illustration of the principle used to estimate head and tail locations of a bee in the first two frames. The bee’s head is shown as the brown circle and the body orientation is shown as the black line attached to the head. The head and tail positions of the bee are labelled as H_1_ and T_1_ for frame 1 and as H_2_ and T_2_ for frame 2. The distances 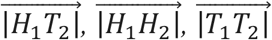 and 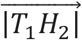 are shown as dashed double arrows in red, blue, brown and magenta, respectively. The motion vector 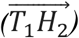 obtained from T_1_ and H_2_ is shown as the dashed red arrow. The vectors 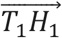 and 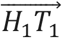 (black arrows) represent the two possible head – tail combinations of the bee in frame 1.

The various scenarios and contingences that are commonly encountered whilst tracking the bees in the cloud are described in the following section, along with methods for dealing with them.

#### Miscellaneous scenarios

##### Occlusions

A common problem that one encounters while tracking multiple bees in video footage is that of occlusion. The first step in handling an occlusion is to detect the event. The presence of an occlusion can be detected by monitoring the areas of the ellipse-fitted images of two bees that are approaching each other. When the image of one bee occludes that of another bee, the ellipses representing the two bees will temporarily be replaced by a single, larger blob. We assume that an occlusion has occurred when this blob contains at least 20 pixels more than each of the individual ellipses.

Once the occlusion frames are detected (and there can be several contiguous occlusion frames), the head and tail positions of each bee in each of the occlusion frames are estimated by extrapolation from their positions in the last two frames prior to the occlusion. (This assumes that, during the occlusion, the image of each bee moves at the same velocity as it did just before the occlusion). At each frame, the predicted (extrapolated) positions of the head and tail of each bee are routinely computed to deal with potential forthcoming occlusions. An occlusion event is detected as an abrupt increase in the area of the fitted ellipse. The extrapolation of the head and tail positions of the two bees involved in the occlusion is continued, frame by frame, over the entire duration of the occlusion. The occlusion is considered complete when the oversized ellipse is replaced by two normal-sized ellipses. In this frame, the extrapolated head and tail positions of the two bees are compared with the ellipses fitted to the two new, unoccluded images that appear after the occlusion, to re-establish the identities of the two bees that reappear from the occlusion, as well as their corresponding head and tail positions. The head and tail positions of the two bees during the occlusion period are then estimated more precisely by linear interpolation, frame by frame, from their positions in the frames immediately before and immediately after the occlusion period. This retrospective linear interpolation provides a better estimate of the head and tail positions of the two bee images during the occlusion period, than the initial linear extrapolation.

##### Tracking bees flying toward the camera’s optical axis

There are instances when a bee moves in such a way as to directly approach the camera, or fly away from it, by moving toward the nodal point of the camera’s lens, or away from it. Under such conditions, the image of the approaching bee will resemble a small circular dot, rather than an ellipse. This situation is assumed to prevail if the ratio of the major to the minor axis of the fitted ellipse is lower than 1.10 but greater than 0.9. In such frames, the centre of the ellipse is taken to represent both the head and the tail positions of the bee’s image.

##### Bees appearing and disappearing from the cameras’ fields of view

When the hive entrance is blocked, the bees which were about to enter the hive will loiter in the vicinity of the hive entrance. After 1 or 2 seconds, some of these bees might choose to leave the arena for a while and come back again to check if they can enter the hive. At the same time, some new returning foraging bees will enter the arena for the first time and will realise that the hive entrance is blocked. So, they will join the cloud and loiter in the arena for some time before leaving the arena. This cyclic process continues until the hive is reopened. So, there will be a continuous flow of incoming and outgoing foraging bees while the bee cloud data is being recorded.

The following procedure is used to record and keep track of the identities bees as they enter and leave the volume of space that is covered by the stereo cameras. The bees that are visible in the first frame of the video from each camera are assigned a unique identification number. These bees are tracked until they leave the arena. Each bee entering the arena during the recording period is assigned a new identification number. (Note that we cannot distinguish between new bees and bees that re-enter the arena: each entering bee is assigned a fresh ID number).

### 3) STEREO MATCHING TECHNIQUES

To obtain 3D reconstructions of the flight trajectories of the freely flying bees, the 2D tracks of individual bees in one camera view have to be matched to the corresponding 2D tracks of the bees in the other camera view. It is therefore crucially important to match the identities of corresponding bees in the two images, in order to obtain correct results. This problem, known as the ‘correspondence problem’ is one of the major challenges in matching stereo images.

We have used two stereo matching techniques - the minimum perpendicular distance method (MPD), and minimum reprojection error method (MRPE), to identify matching bee images in the two camera views. The details of the two methods are discussed below.

#### a) Minimum perpendicular distance method

The minimum perpendicular distance method was first used in a 3D particle tracking velocimetry algorithm^41^ as an intermediate step for solving the problem of correspondence matching. Later, this basic idea was modified and applied to solve the stereo matching problem in a study involving midge swarms^18^ and bird flocking events (jawdacks and rooks)^42^ using three cameras. Here, we use this technique to solve the ‘correspondence problem’ with just two cameras. The concepts and steps involved in implementing this algorithm are discussed below.

Consider a bee at a location *M*_*1*_ in 3D space, which is imaged at points *m*_*11*_ and *m*_*12*_ in the image planes of cameras 1 & 2, respectively, as shown in Fig. 3. (The points correspond to the centroids of the ellipses fitted to each image.) We construct a back-projected ray vector 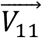 which emanates from the nodal point of Camera 1 (*c*_*1*_) and passes through the point *m*_*11*_ in the camera’s image plane. Similarly, we construct a back-projected ray vector 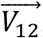 which emanates from the nodal point of Camera 2 (*c*_*2*_) and passes through the point *m*_*12*_ in its image plane. If the images at *m*_*11*_ and *m*_*12*_ are from the same bee, the back-projected vectors 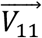 and 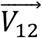 should intersect at a location in 3D space (say, *M*_*1*_). However, due to the presence of noise and imperfect camera calibration, the vectors will not intersect: they will be separated by a small perpendicular distance Δ*d*_*11*_, as shown in Fig. 3. If the images in Cameras 1 and 2 are from different bees, as depicted by *m*_*11*_ and *m*_*12*_, the perpendicular distance Δ*d*_*12*_ between the two back-projected rays will, in general, be greater than Δ*d*_*11*_. Correct correspondence can therefore be achieved by determining, for a given bee image in Camera 1, the bee image in Camera 2 that produces the lowest perpendicular distance between the back-projected rays. This is the ‘Minimum Perpendicular Distance’ method that we use to solve the ‘correspondence problem’. To implement this method, we need to compute the perpendicular distance between the back-projected rays from Cameras 1 and 2. This procedure is described below.

**Figure 3.**
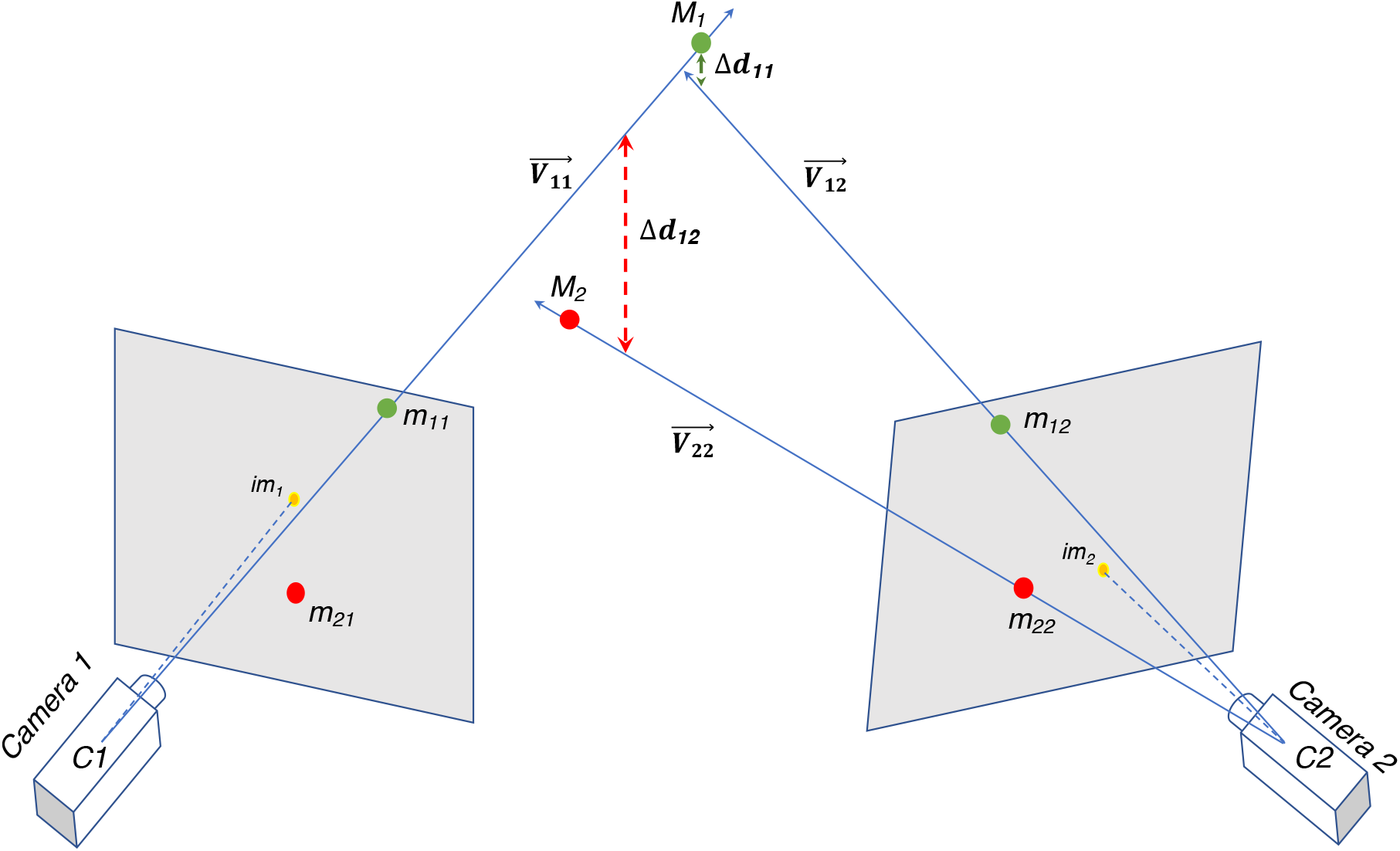
Schematic illustration of the concept behind the minimum perpendicular distance method showing the shortest distances – ‘Δd_11_’ and ‘Δd_12_’, when a matched or a mismatched pair of images are stereo-matched. C1 and C2 denote the nodal points of Camera 1 and Camera 2, respectively. The image positions m_11_ and m_12_ are those of a correctly matched pair of images of a bee at location M_1_. This results in a relatively small perpendicular distance Δd_11_, as shown by the vertical dashed green line. On the other hand, a pair of mismatched images (such as m_11_ in Camera 1 of the bee at M_1_, and m_22_ in Camera 2 of a different bee at location M_2_) result in a larger perpendicular distance Δd_12,_ as shown by the dashed red vertical line.

The perpendicular distance, in 3D, between any two ray vectors can be computed as follows: We define 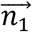 and 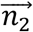 to be unit vectors representing the directions of the back-projected vectors 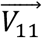 and 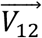, respectively (see Fig. S1, Supplementary File 3). These can be readily computed, because 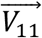 and 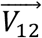 are known. (In fact, 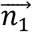 and 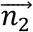 do not need to be unit vectors: they can be of any length, and their lengths need not be equal). The direction of the unit perpendicular distance vector 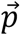 is then given by the cross product of *n*_1_ and *n*_2_:

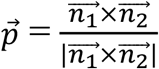

*C*1 and *C*2 are arbitrary points on the back projected vectors 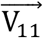and 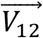, respectively. Here, we have chosen *C*1 and *C*2 to be the nodal points of Cameras 1 and 2, respectively.

Then, as shown in the derivation in Fig. S1, the perpendicular distance *P* is given by the projection of 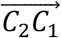 on 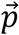, which is obtained from the magnitude of their dot product:

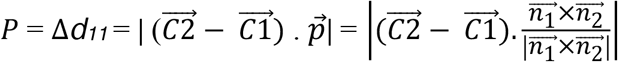

Our perpendicular distance-based stereo-matching algorithm processes the image sequences from the two cameras as follows:

1. The 2D trajectories of the images of the bees captured in the video from Camera 1 are considered to be the reference tracks. The algorithm compares each reference track from Camera 1 with each of the tracks in the video from Camera 2.
2. Before considering a 2D track from Camera 2 for further processing, the program checks if this track has video frames that overlap with the frames of the reference bee’s 2D track. If there are no overlapping frames, it means that the tracks in the two cameras did not overlap at all in time, which implies that the two tracks in question arose from different bees. This is the first step in the procedure for eliminating mismatched tracks in Camera 2.
3. Next, the algorithm performs a comparison between each 2D reference track from Camera 1, and all of the surviving 2D tracks from Camera 2. For each coinciding frame between a reference track in Camera 1 and all of the surviving tracks in Camera 2, we compute the perpendicular distance between the ray vectors emanating from Camera 1 and Camera 2. For example, consider a comparison of the first frame of reference track #1 from Camera 1with 80 different surviving tracks, in the corresponding frame from Camera 2. This comparison will provide 80 perpendicular distance values for this frame. We repeat this step for each of the overlapping frames, and compute a mean perpendicular distance for each of the 80 candidate tracks in Camera 2. This yields a total of 80 mean perpendicular distances. Three valuable inferences can be drawn from this array of 80 mean perpendicular distances: (a) We consider the value of the minimum mean perpendicular distance. If this value is lower than 10 mm, then the corresponding track in Camera 2 is the one that matches the reference track #1 from Camera 1. (b) If the first and second lowest mean perpendicular distances (arising from two candidate tracks in Camera 2) are both less than 10 mm, and if the difference between them is less than 6 mm, then the two well matching tracks arise from the same bee appearing twice in Camera 2’s field of view, during different time windows. In this case, we treat the mean perpendicular distance values obtained for the two candidate tracks as originating from the same bee, and we use the average of these two values to represent the appropriate mean perpendicular distance for that particular combination of tracks. (c) If the value of the mean perpendicular distance is greater than 10 mm for all of the 80 candidate tracks, then the bee corresponding to reference track #1 from Camera1 was never in the field of view of Camera 2. This enables the identification of the bee track in the video from Camera 2 that best matches the track of reference track #1 as well as the exclusion of bees that are in the field of view of Camera 1, but are not visible in Camera 2. [Note that the criterion distance (10 mm) is selected and fine-tuned by the user, based on the density of bees in the cloud].
4. On completion of steps (1-3) we have, from the 80 tracks in Camera 2, the candidate track that provides the best stereo match for reference track #1 from Camera 1. This candidate track in Camera 2 is taken to be the one that was produced by the bee that generated reference track #1 in Camera 1. That is, these two tracks are labelled as constituting a stereo pair generated by bee #1.
5. This procedure is now repeated by treating each of the remaining tracks in Camera 1, in turn, as the reference bee, and finding its corresponding stereo-matched track in Camera 2. We note that, with each successive reference track that is considered, the number of Camera 2 tracks for comparison decreases by one, because of the progressively increasing number of tracks that have already been stereo-paired.

#### b) Minimum reprojection error method

Another approach for solving the stereo correspondence problem is to use Matlab’s ‘triangulate’ command^43^. The ‘triangulate’ function, in general, returns the 3D location of a matching/non-matching pair of 2D image points from any two stereo images. But it does not tell us whether it has triangulated a correct pair of image points or not. However, we can infer this information by looking at the reprojection error (RPE) parameter returned by the ‘triangulate’ function in Matlab.

Basically, the reprojection error in each camera’s image plane is the distance (in image pixels) between the centroid of the image of a bee, and the centroid of a back-projected image of the 3D position of the bee, as estimated by Matlab’s stereo algorithm. Matlab delivers the value of the reprojection error registered in the two camera images.

In general, the RPE is low when the images in the two cameras are matched (i.e. when they correspond to the same bee) and high when the images are not matched (i.e. when they correspond to different bees). We use a RPE threshold of 10 pixels: images in the two cameras are considered to belong to the same bee if their RPE is less than or equal to 10 pixels, and to different bees if their RPE is greater than 10 pixels. This error threshold was selected after inspecting the RPE values for a large number of arbitrarily paired bee images.

The procedure for discovering matching pairs of trajectories in the two camera images is the same as that described for the minimum perpendicular distance method, except that in this case the minimum perpendicular distance criterion is replaced by the minimum reprojection error criterion. For any given reference bee image in Camera 1, if the mean RPEs for all of the 80 candidate bee images in Camera 2 are greater than 10 pixels, then we conclude that there is no corresponding bee image in Camera 2. And if the mean RPEs for any two bee image pairs are lower than 6 pixels, we assume that they both represent matching image pairs of the same bee, and that this bee has entered the field of view of Camera 2 twice.

To our knowledge, none of the previous stereo tracking studies^24,35,42,44^ have evaluated the accuracy of their stereo-matching algorithms. We conducted a control experiment to assess the accuracy of the two stereo-matching techniques described above. This control experiment is described in the following section.

#### c) 3D Reconstruction of bee trajectories

When a reference bee is correctly stereomatched with its counterpart in Camera 2, then the unique ID number of that particular reference bee is saved. This process is repeated for each reference bee, to find all of the stereomatched pairs in the data. Using the obtained unique ID numbers, for each stereo-matched pair of bee image trajectories the respective 2D head and tail positions (in image coordinates) are passed to a triangulation routine which generates the 3D head and tail trajectories of that particular bee. This procedure is repeated for all of the matched image tracks to obtain the 3D head and tail locations of each bee in every frame and to reconstruct 3D trajectory of each bee.

### 4) CONTROL EXPERIMENT

The control experiment used the same camera system, but a different arena. The experiment used twelve balls, of different sizes and colours, which were individually identifiable in each camera by visual inspection. A 1.96m x 1.56m sheet of white corflute was laid on the ground. The positions and orientations of the two cameras were arranged to ensure that each camera’s field of view fully covered the corflute sheet. The 12 balls were dropped from a height on the corflute sheet and were filmed by the two video cameras at 25 frames/sec as they bounced. The white background of the corflute sheet provided a high-contrast background against which the individual balls could be detected and identified accurately. Therefore, it was possible to match the corresponding images of each ball in the two camera images manually, with 100% accuracy, frame by frame. This process of manual matching provided the ground truth information, against which the two automated stereo-matching methods described above could be compared. Before recording the data, the usual stereo camera calibration procedure was performed to obtain the intrinsic and extrinsic parameters of the two cameras.

Next, a few random image pairs were extracted from the captured video data. One such image pair is shown in Fig. 4, which depicts 10 of the 12 balls used in the experiment. The centre of mass (CoM) of each ball in the scene was manually digitised using the ‘ImageJ’ tool, to obtain its pixel coordinates in the 2D image. The corresponding images of each ball in the two views were matched manually, as each ball could be unambiguously identified in the two images, based on its colour and size. This manual matching provided the ground truth for the control experiment, which was used to validate and compare from the accuracy of the two automated stereomatching algorithms. As per our usual convention, the ball images in Camera 1 were considered as reference images, and the two stereomatching algorithms were run to select the corresponding images in Camera 2, for each reference ball in Camera 1.

**Figure 4.**
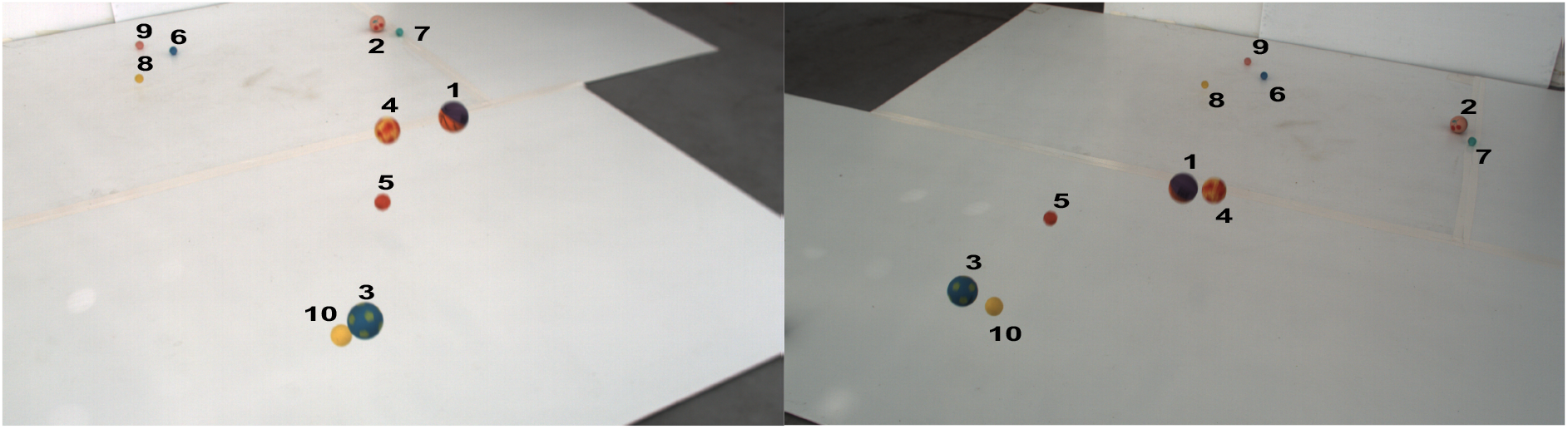
A stereo-pair captured by the camera system during the control experiment. The manually matched images of each ball in the two images are indicated by the numbers. This particular image pair contains only 10 of the 12 balls used in the experiment, as the other two balls had left the fields of view of both cameras by then.

## TECHNICAL VALIDATION

### 1) VALIDATING 2D TRAJECTORIES

As we do not have any ground truth information on the 3D trajectories of the bees flying in the cloud, it was not possible to perform a direct quantitative evaluation of the accuracy of the trajectories generated by our algorithms. However, the developed algorithms can be used to identify and eliminate various tracking errors and mismatches. It is also possible to validate the algorithms by using known parameters such as the body length and the expected turning rates of honeybees, as we show below.

#### (a) Body orientation and length

The orientation of the bee’s body, as obtained from the computed estimates of the 3D positions of the head and the tail, can be used to test the validity of the 3D reconstructions of the bees’ trajectories. Given the high video frame rate (200/s), one would not expect the orientation of the bee’s body to change substantially between successive frames. A large change in the computed orientation between frames would occur only if (a) the head and tail positions of a bee’s image have been erroneously swapped, or (b) if the images in the two frames do not represent the same bee, due misidentification. Fig. 5a shows the magnitude of the maximum change in body orientation (Δ*θ*), between successive frames for each bee. It is clear that the (Δ*θ*) values are consistently low, with the mean being 8.0 deg, and 90% of the values being lower than 6.2 deg. This confirms that there has been no misregistration of orientation, or misidentification of bees in the process of reconstructing the 3D trajectories, thus proving one form of validation.

**Figure 5.**
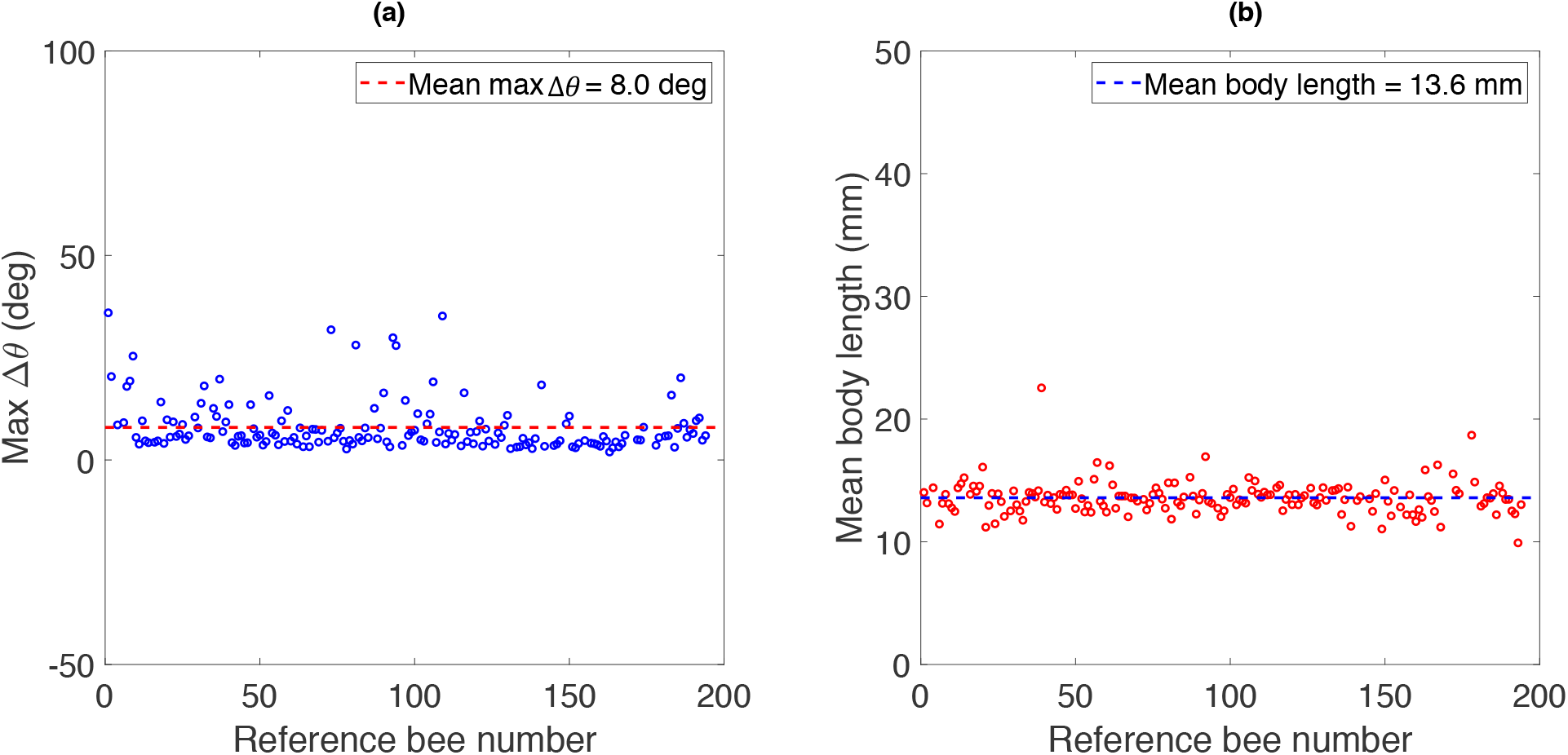
(a) Maximum absolute value of the frame-to-frame change in the body orientation of each bee. (b) Mean body length of each bee, averaged over its entire flight trajectory.

Another way to check the validity of the reconstructed trajectories is to examine the lengths of the bodies of the bees, from the computed 3D positions of the head and the tail. The mean body length, averaged over all bees, is 13.6 mm ± 1.32 mm (SD), (see Fig. 5b), which agrees very well with the known average body length of around 12-15 mm^45^ for the honeybee (*Apis mellifera*).

#### (b) Manual validation

Once a bee was tracked, we plotted its 2D head and tail positions, and body orientations, over its entire flight. We performed a high-level check to see if the tracked 2D trajectory resembled the original flight path of the bee by visually comparing the tracked 2D trajectory with the flight path of the bee in the raw footage. We also checked for any visually obvious abnormal changes in the direction of flight of each bee’s 2D trajectory. The 2D trajectories were then overlaid on their respective camera views to verify that the tracked head and tail positions matched the corresponding bee images in each camera view. Two such video excerpts where the generated 2D trajectories are overlaid on the respective camera views are given in Supplementary File 1.

While reconstructing the 3D trajectories of the bees, we noticed that the trajectory would occasionally go outside the boundaries of the arena. One possible reason for this erroneous 3D track is misidentification of the bee (i.e., an identity swap) in one of the camera views. This can occur particularly when there is an image of another bee near the image of the tracked bee, which is also flying in approximately the same direction. Fortunately, our matching algorithms provide a means for detecting such identity swaps. When the tracking of a bee in one of the camera views is abruptly switched to a different (incorrect) bee, this shows up as an abrupt and large increase in the MPD and the MRPE. Apart from obtaining a high value of MPD and MRPE, at times, one might also observe a sharp discontinuity in the reconstructed 3D trajectory, possibly taking it outside the arena boundary.

### 2) RESULTS OF THE STEREO-MATCHING TECHNIQUES, AND OF THEIR APPLICATION AS A TRAJECTORY VALIDATING PROCEDURE

In this section, we first show the results obtained from the two stereo-matching programs when they are applied to the 2D trajectories of the bees obtained from the two camera views. We also describe how we verify the validity of the results. Finally, we describe the results of the control experiment for evaluating the accuracy of the stereo-matching techniques developed and used in this study.

As described in the ‘Methods’ section we compute, for each reference bee in Camera 1, the minimum mean perpendicular distance (MMPD) for each of the bee trajectories in Camera 2 to obtain a set of mean perpendicular distances (MPDs). The number of MPDs obtained will vary from one reference bee to the next, because this will depend on the number of frame synchronised bees that are present in Camera 2. The pair that yields the lowest MPD (i.e. the MMPD) is considered to represent the matching pair. The results are shown in Fig. 6, which presents the results obtained for bees 1-100 (Fig. 6a) and for bees 101-200 (Fig. 6b), in each of which the image sequences of 100 bees were matched. The MMPD value obtained for each reference bee is shown as a blue dot, and the flanking blue bars represent the standard deviation of the perpendicular distance (PD) measured across all of the frames in which this bee was present in both camera views. The overall mean value of the MMPD, averaged across all the reference bees, was found to be 2.06 mm. The second-lowest MPD values are shown in green, with the green dots and vertical lines representing the mean and standard deviation of the PD values. The overall mean value of the second lowest MPD, averaged across all the reference bees, was found to be 57.84 mm, which is significantly greater than the lowest MPD (*p =*1.84 × 10^−57^, *one sided t-test*). The magenta dots show the averages of the MPD values (the third-lowest value to the highest value) for the remaining bees, and the flanking bars show the standard deviations of these values. It is clear that the MPD values obtained for the remaining bees are substantially (and significantly) higher than that obtained for the bee which generates the lowest MMPD, which is the matching bee (*p =* 3.01 × 10^−106^, *one sided t-test*). For each reference bee, the corresponding matching bee always yields a MMPD that is lower than 10 mm. For all non-matching bees, the MMPD is considerably greater than 10 mm. These observations confirm the high reliability with which the MMPD algorithm selects correctly matching bees in the two camera views.

**Figure 6.**
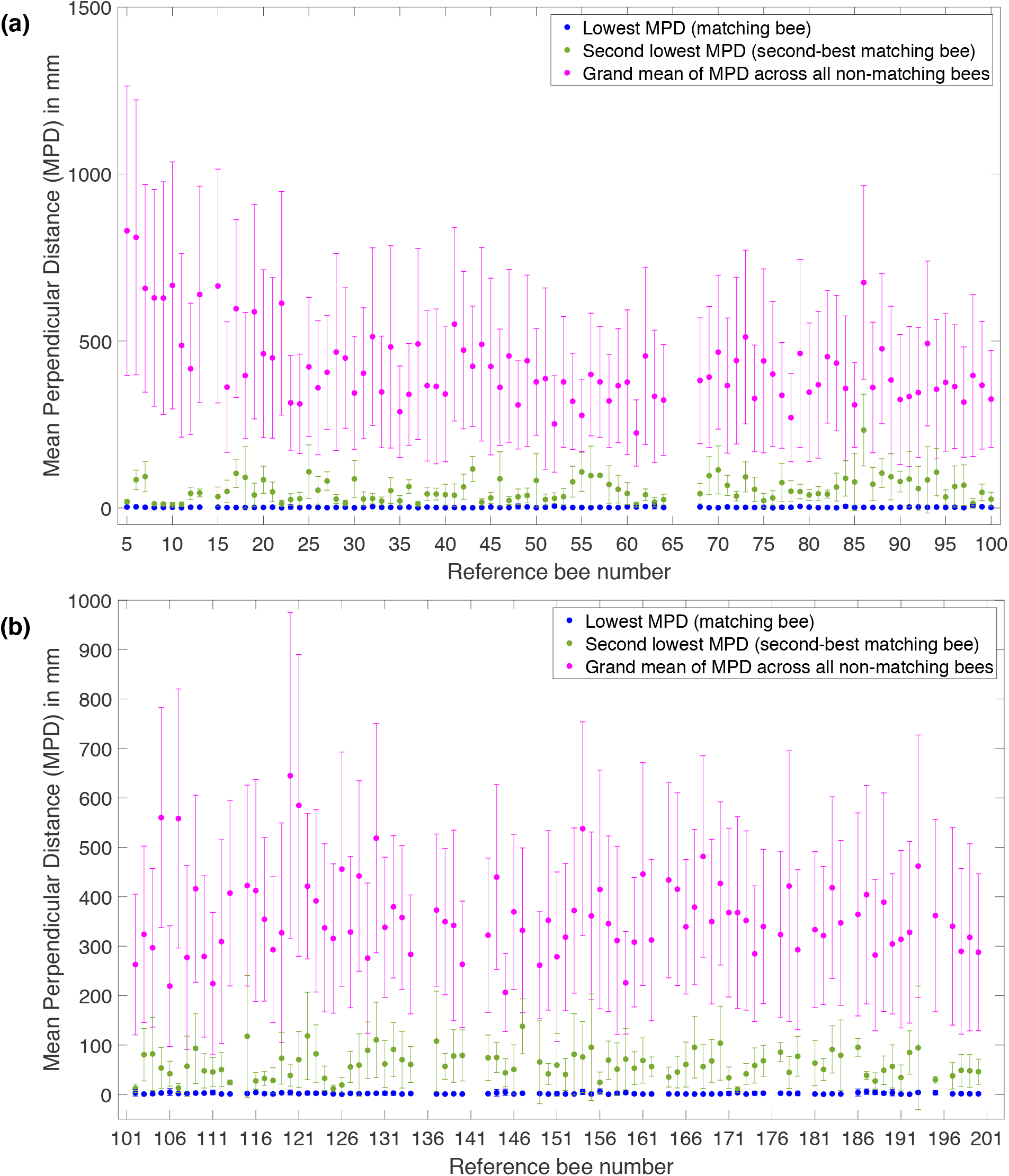
The Mean Perpendicular Distances (MPDs) computed for the first set of 100 reference bees (bees # 1-100) **(a)**, and for a second set of 100 reference bees (bees # 101-200) **(b**). Details in text.

Fig. 7 shows an analysis similar to that in Fig. 6 but compares the reprojection errors (RPEs), rather than the perpendicular distances, for the same 200 reference bees. The lowest mean RPE (blue dots), averaged across all reference bees, was 3.44 pixels. The second lowest mean RPE (green dots), averaged across all reference bees, is 31.35 pixels, which is significantly greater than the lowest mean reprojection error (blue dots) (*p =* 5.40 × 10^−62^, *one sided t-test*). The magenta dots show the averages of the RPE values (the third-lowest value to the highest value) for the remaining bees, and the flanking bars show the standard deviations of these values. It is clear that the RPE values obtained for the remaining bees are substantially (and significantly) higher than that obtained for the bee which generates the lowest RPE, which is the matching bee (*p =* 8.68 × 10^−113^, *one sided t-test*). For each reference bee, the corresponding matching bee always yields a mean RPE that is lower than 10 pixels. For all non-matching bees, the mean RPE is considerably greater than 10 pixels. These observations demonstrate the good accuracy with which the RPE algorithm selects matching bees in the two camera views. However, as we shall show below, the MMPD algorithm is slightly more reliable than the MRPE algorithm, on several counts.

**Figure 7.**
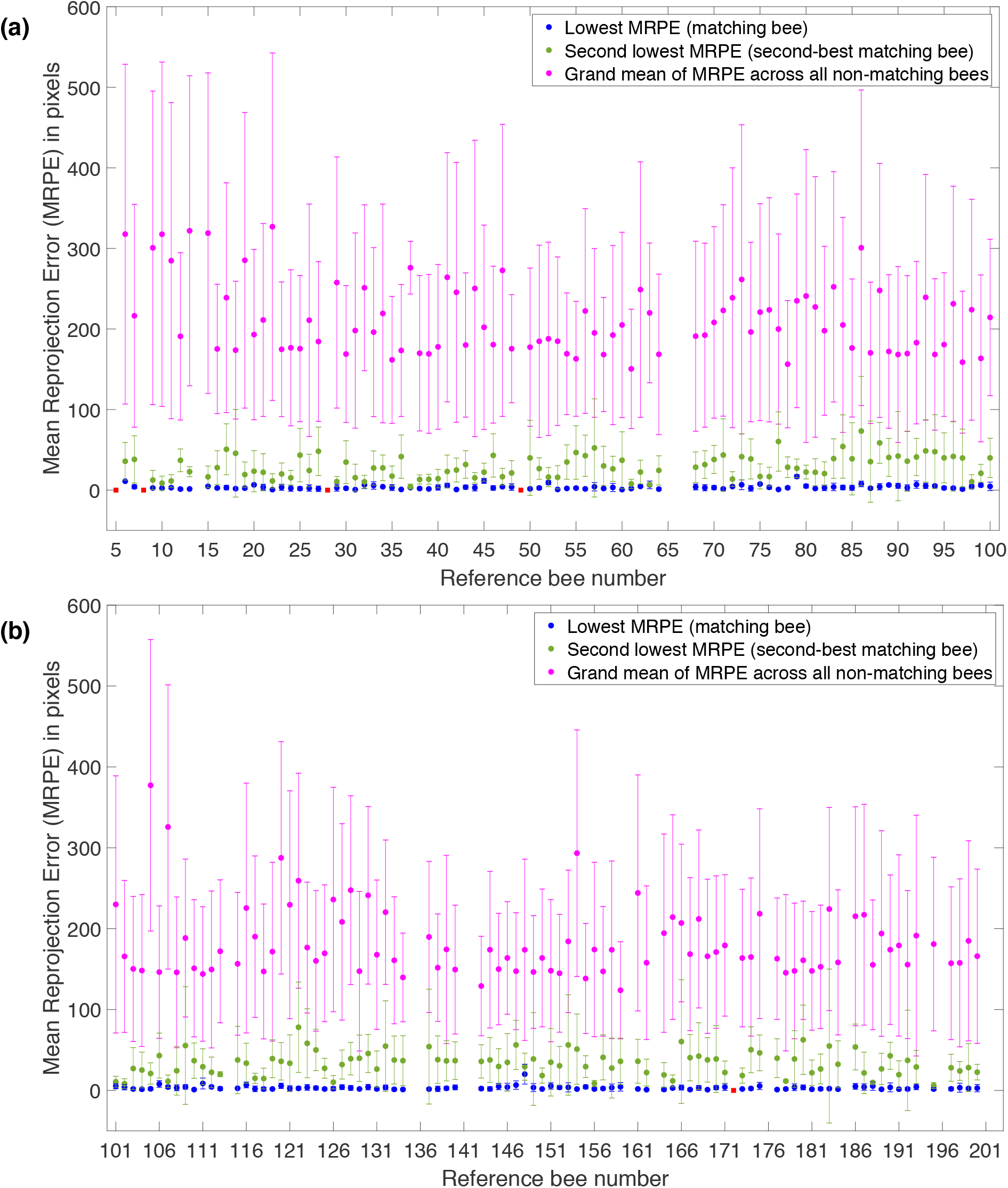
Mean Reprojection Error (MRPEs) computed for the first set of 100 reference bees (bees # 1-100) **(a)**, and for a second set of 100 reference bees (bees # 101-200) **(b)**. The red dots represent instances in which the MMRPE yields the incorrect match, but the MMPD yields the correct match. The other symbols have the same meanings as in Fig. 6. Details in text.

Firstly, we compared the accuracies of the two algorithms in selecting matching image pairs by examining the best-matching pair and the second best-matching pair. In some instances both algorithms yielded two pairs that met the respective matching criteria (MMPD < 9 mm, and MMRPE < 7 pixels), but the pair selected by the MMPD algorithm was different from the pair selected by the MMRPE algorithm. These instances were rare – they occurred only in reference bees 5, 8, 28, 49 and 172, and are indicated by the red dots in Fig. 7. For these instances we inspected the videos from the two cameras visually to determine the correctly matching pair, by examining (a) the similarity of the direction of movement of the images of the candidate bees in the two camera views; (b) synchronised turning of the bee images in the two camera views^46^; and (c) similarities in the points where the candidate bees in the two views appeared to move outside the arena boundary. We found that, in all five instances, the MMPD algorithm determined the correct matching pair, while the MMRPE method selected the wrong pair. Thus, the MMPD method is 100% accurate in selecting the correctly matching pair for all of the reference bees. The MMRPE method is nearly as accurate, displaying an error rate of 5 in 200 examples, or 2.5%.

With both methods, the accuracy improves with the duration of the trajectory (the number of frames), because mismatches occur only in a small percentage of the frames. Matching errors are more likely to occur with short trajectories (e.g. bee # 5,8) because mismatches in even a small number of frames would have a significant effect on the computed overall MMPD and MMPRE. Nevertheless, even with the shorter trajectories, we find that the MMPD method consistently yields the correct matches.

As mentioned earlier, the stereo-matching algorithm can be used to detect erroneous identity (ID) swaps that occur during the tracking process. This could happen, for example, when a tracked bee abruptly disappears from the field of view of one camera, and its subsequent track is replaced by that of a neighbouring bee that remains in the field of view. In order to demonstrate the ability of the algorithm to detect such errors, we created 2D trajectories of bees with their identities deliberately swapped with other bees. We did this by creating a trajectory in which the 2D head and tail positions of a particular reference bee were used up to the halfway point in its flight. Beyond this point, the 2D head and tail coordinates of a different bee (which was also flying at the same time in the arena), were appended to the trajectory of the reference bee previously considered. This is one way of emulating an accidental identity swap scenario for testing purposes. The usual procedure of minimum distance and reprojection error-based stereo-matching techniques were then applied to these manipulated trajectories. The results from the two stereo-matching techniques for five correct and five ID swapped trajectories are shown in Table 1.

**Table 1.** Lowest mean minimum distance and lowest mean RPE values obtained from 5 reference bees, when their trajectories contained (left) or did not contain (right) deliberately introduced ID swap errors.

| Reference bee # | Trajectories with ID swaps |  | Trajectories without ID swaps |  |
| --- | --- | --- | --- | --- |
|  | Lowest MPD (mm) | Lowest MRPE (pixels) | Lowest MPD (mm) | Lowest MRPE (pixels) |
| 7 | 326.6 | 69.7 | 2.1 | 4.2 |
| 50 | 47.1 | 24.3 | 0.9 | 1.6 |
| 30 | 72.4 | 21.0 | 0.6 | 2.2 |
| 32 | 135.8 | 65.4 | 4.1 | 6.6 |
| 12 | 256.3 | 147.7 | 2.2 | 1.3 |

It is evident from this table that the lowest MPD and MRE values are considerably larger when ID swaps are introduced into the reference trajectories, than when they are not. Therefore, the two stereo-matching algorithms are capable of reliably detecting when the 2D trajectory of a particular bee in the video sequence from Camera 1 is erroneously swapped, midstream, with the 2D trajectory of another bee.

In addition to the validation procedures discussed above, we performed a control experiment to assess the accuracy of the two stereo-matching techniques. The details of the control experiment are given in the Methods Section, and the results are discussed below.

A random image pair was selected from the video footage of the control experiment. The 2D coordinates of each ball were digitised (manually) in the two camera views in a particular order for better visualisation of the results. The balls in the two views were digitised in the following sequence: (1) orange-violet basketball, (2) red-blue basketball, (3) pink-red ball, (4) blue football, (5) yellow football, (6) small red ball, (7) small blue ball, (8) small green ball, (9) small yellow ball, (10) small pink ball, (11) small light pink ball, and (12) yellow plastic ball. It was possible to visually match images of corresponding balls in the two images manually with 100% accuracy, as each ball could be uniquely identified from its colour and size. This provided the ground truth for assessing the accuracies with which the MPD and MRPE algorithms selected the correctly matched image pairs.

Fig. 8a shows the results obtained by applying the PD and RPE algorithms in a randomly selected (time-synchronized) pair of camera images. The numbers in the pairing matrix show the computed PD and RPE values for each visible candidate pair. It is clear that both the PD and the RPE values are lowest for the correctly matched pairs, highlighted by the maroon cells along the diagonal of the pairing matrix. This demonstrates that both algorithms are capable of selecting all of the correctly matched image pairs - except in those cases in which a ball is not present in one (or both) camera images, as signified by the ‘NA’ symbols.

**Figure 8.**
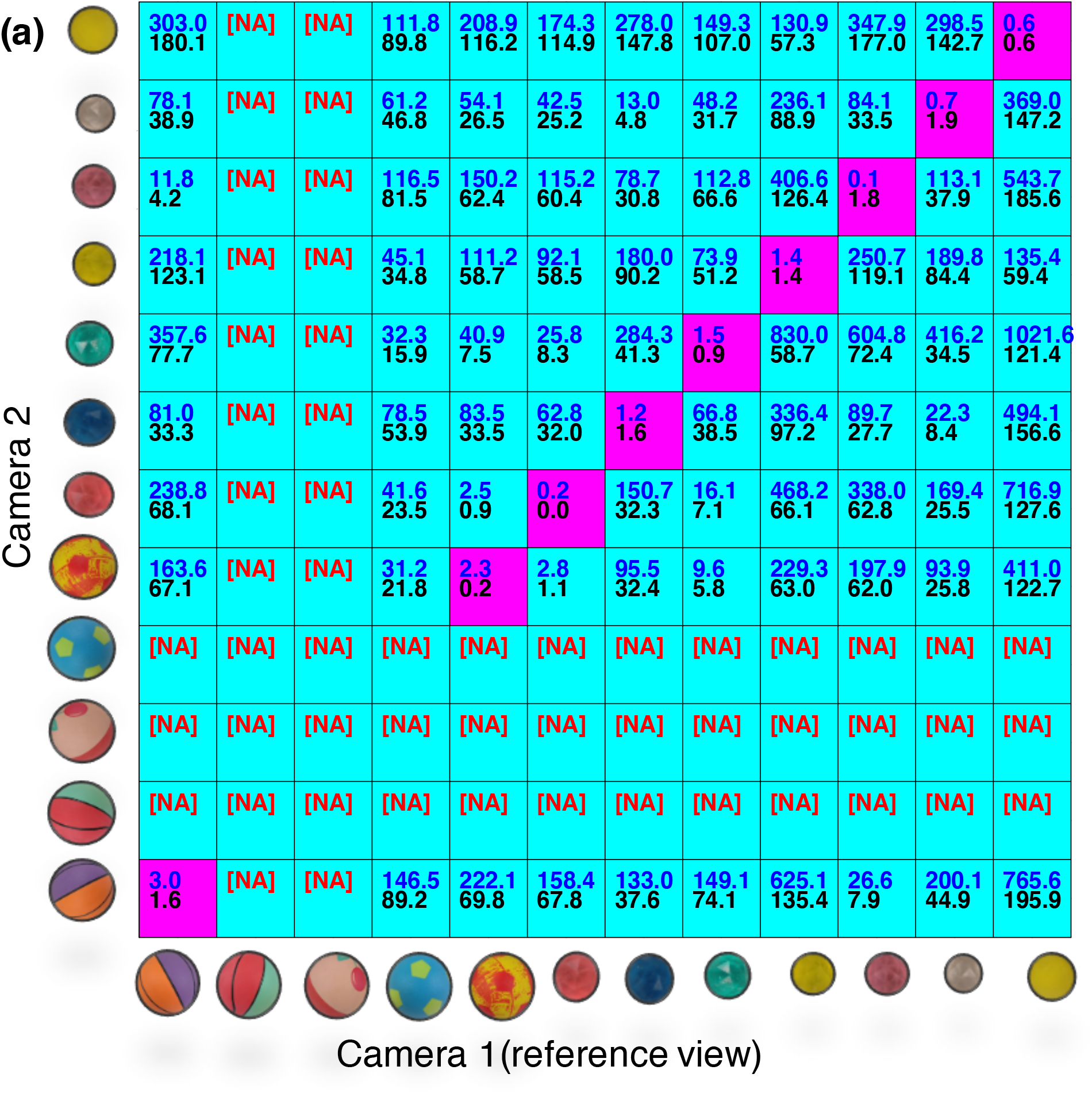
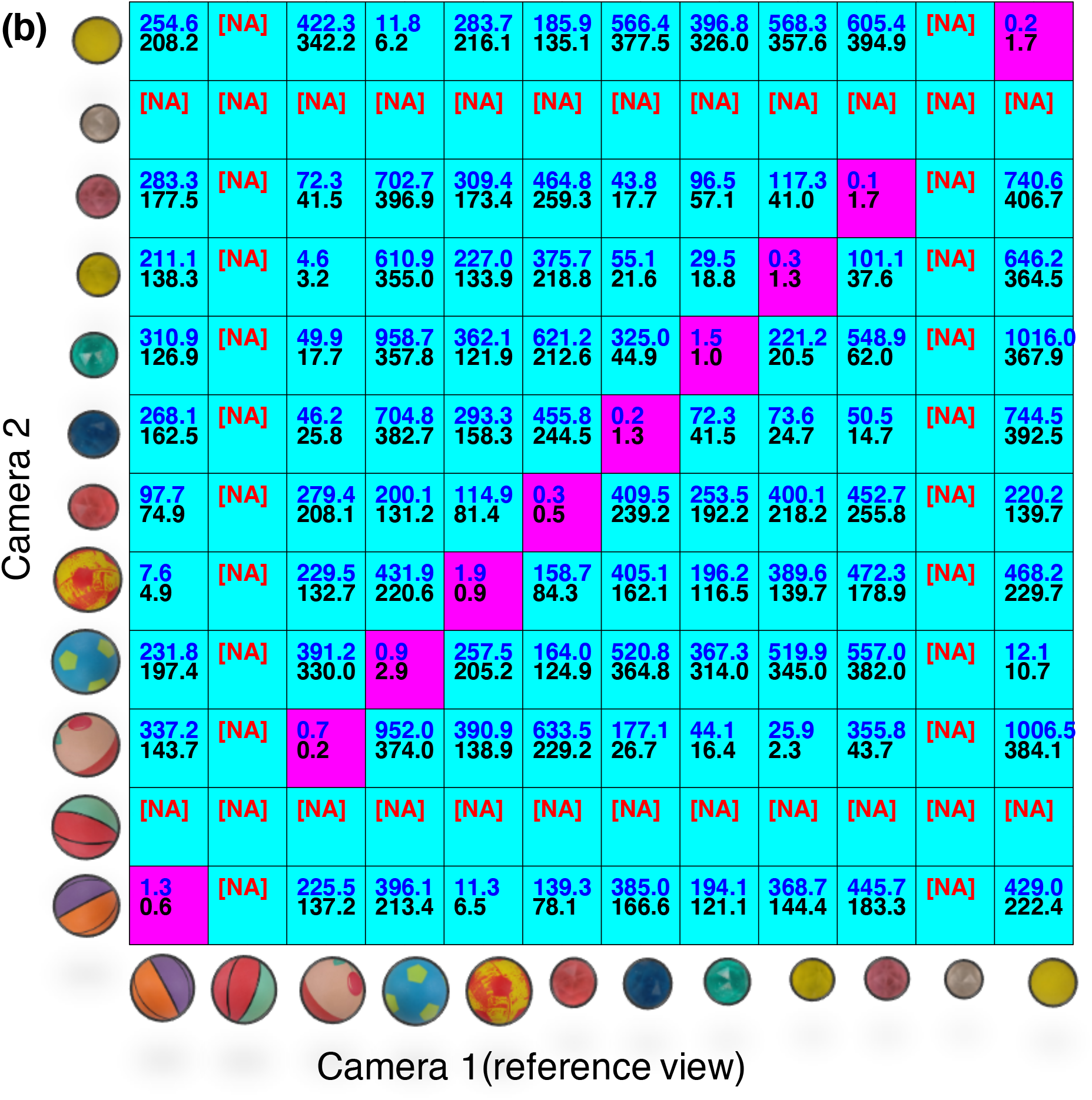
Results of the control experiment, showing a comparison of the perpendicular distance (PD) and the reprojection error (RPE) values computed from two randomly selected image pairs (a, b), for all possible candidate ball pairs. The PD and RPE values are shown in blue and black, respectively. The lowest PD and RPE values are obtained for the correctly matched pairs, as shown by the cells highlighted in maroon, which lie along the diagonal of the pairing matrix. **NA** implies that the particular ball was not available in that camera view.

Fig. 8b shows the results obtained from another randomly selected image pair. Again, both algorithms are able to match all visible pairs with 100% accuracy. This control experiment provides another validation of the two matching algorithms developed and examined in this study.

The flight trajectories of 15 different bees that were visible the first 300 frames of bee cloud data 1 are shown in Fig. 9. It should be noted that there were as many as 13 other bees flying in the same time window, but all of the bees were not included in the plot to ensure clear visualization of the individual trajectories. In total, there are 382 individual 3D bee trajectories in the two bee cloud datasets, comprising 121544 head and tail positions (dataset-1: 70771 head & tail positions, dataset-2: 50773 head and tail positions). The Supplementary File 2 (movie S3 & S4) shows the trajectories of all bees flying in the two sets of bee cloud data that were filmed, digitised and analysed in this study.

**Figure 9.**
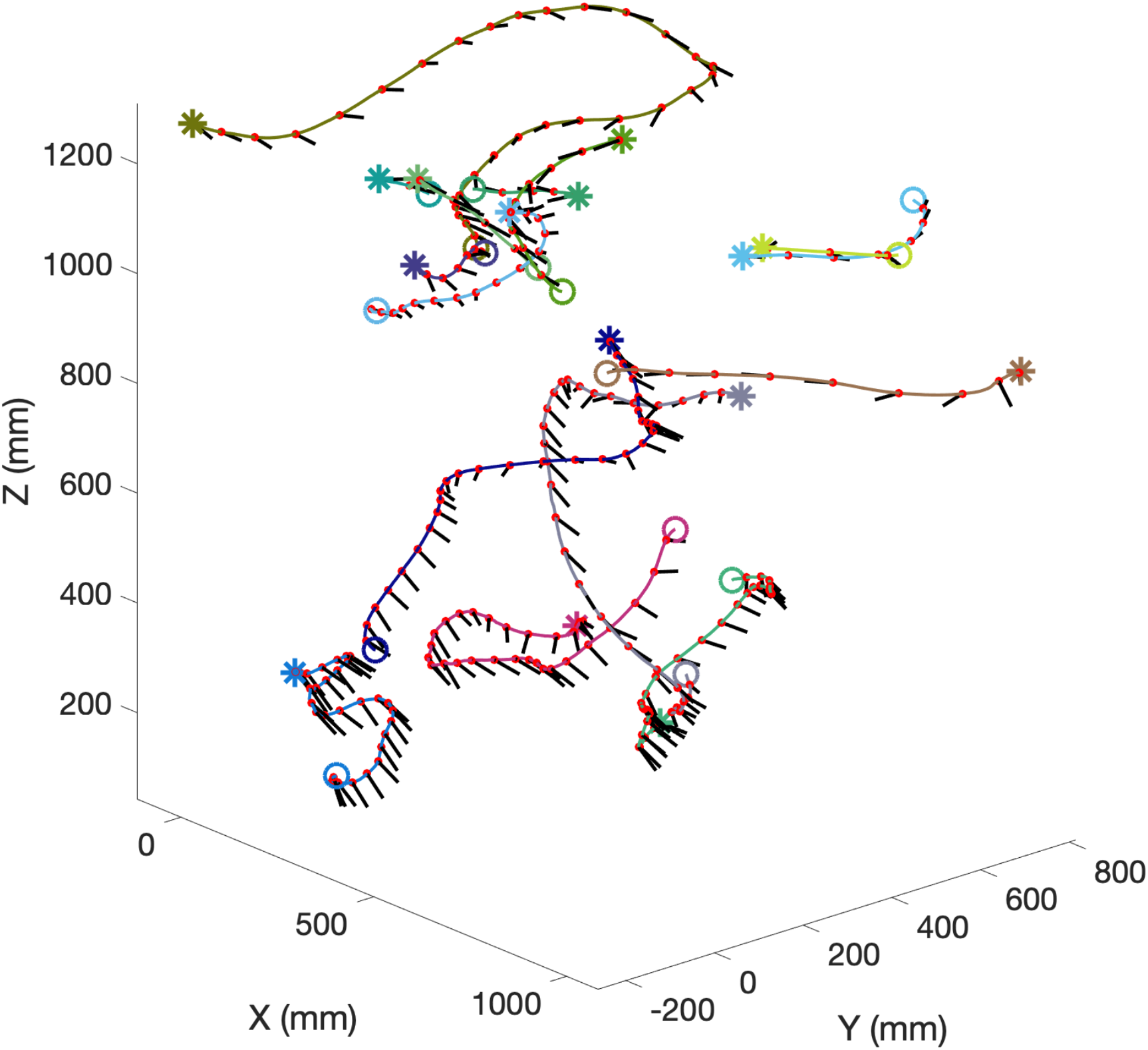
3D reconstructed trajectories of 15 different bees that were imaged during the first 1.5 seconds (300 frames) of the recorded bee cloud data 1. The flight path of each bee is shown in a different colour. The starting point of each trajectory is indicated by a hollow circle and end point by an asterisk. The head position and body orientation of each bee are shown as a red dot and a black line, respectively, at 10 frame intervals.

## DATA RECORDS

The dataset (Data Citation 1) consists of head and tail positions of bees filmed in two different bee cloud events. The first and second bee cloud data were recorded for 7.9 & 7 seconds, respectively. The head and tail positions of each bee across the frame sequences are provided in separate text (.txt) files. The first column is the unique ID number of each reference bee. The second column specifies the frame number in the video sequence. The third, fourth and fifth columns in each file provide the *x, y* and *z* coordinates of the head (or tail) in mm with respect to the coordinate system of the arena. The sixth column contains a flag variable (‘H’ or ‘T’), which indicates whether the *x, y* and *z* coordinates represent the head or the tail position of the bee. Finally, the seventh column indicates whether the data in the file correspond to the first or the second bee cloud.

We have named each file according to the format ‘CLOUD_EVENT_*_BEE_#_HEAD_POS.txt’ and ‘CLOUD_EVENT_*_ BEE_#_TAIL_POS.txt’, where ‘#’ is the unique ID number of each bee in the data and ‘*’ is the bee cloud event number. Thus, the file name also identifies whether it carries data on head positions or tail positions.

## Code availability

The code for tracking and stereomatching is available at https://github.com/mahadeesh/Bee-tracking-and-stereomatching-files.

## Data Citations

1. https://figshare.com/s/bc52d39dd87d9dda54d5

## Acknowledgements

Our special thanks to Mr. Peter Ryan, for the care of the bee colony used in this experiment. This study was supported by a UQ International student scholarship, a Boeing top-up scholarship awarded to M.M., and partly by Australian Discovery Research Grant DP 140100896 to M.V.S.

## Author Contributions

M.M. and M.V.S. designed the experiments. M.M. analysed the data. M.M. and M.V.S. wrote the manuscript.

