## Supplementary Information for "Reconstruction of Three-Dimensional Trajectories of Honeybees Flying in High-Density Aerial Environments"

*SUPPLEMENTARY FILE 1*

***Manual validation of the 2D trajectories in each camera view***

A high-level manual verification was performed on the bee trajectories in each camera view to make sure there are no visible tracking errors in the data such as a switch in head and tail positions, identity swap issues. This was done by visually inspecting each trajectory, once it has been tracked by our tracking program. Finally, once the bees in each camera view are tracked, we overlaid its head and tail position on to its respective camera view. This procedure was repeated for each camera view in the two datasets and the resulting videos are shown below.

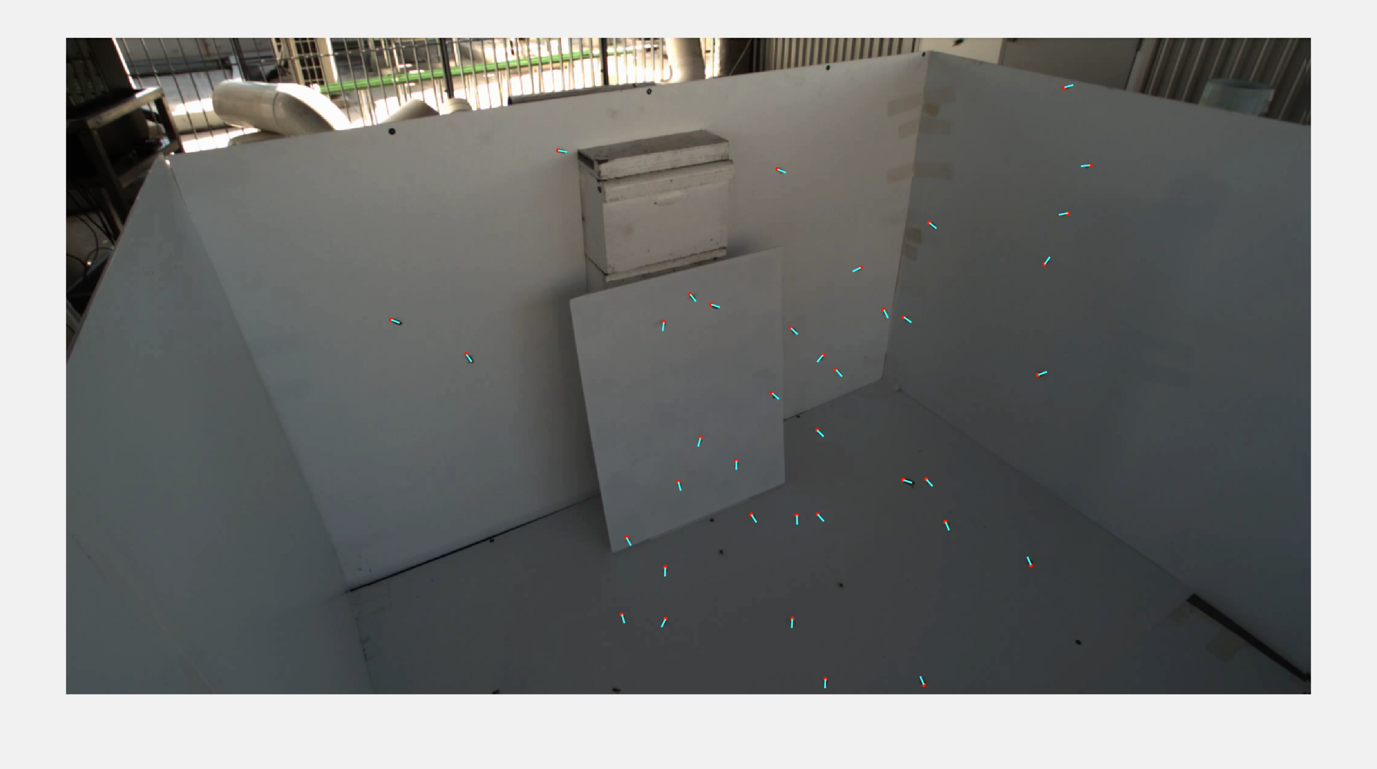
(S1)

(S2)
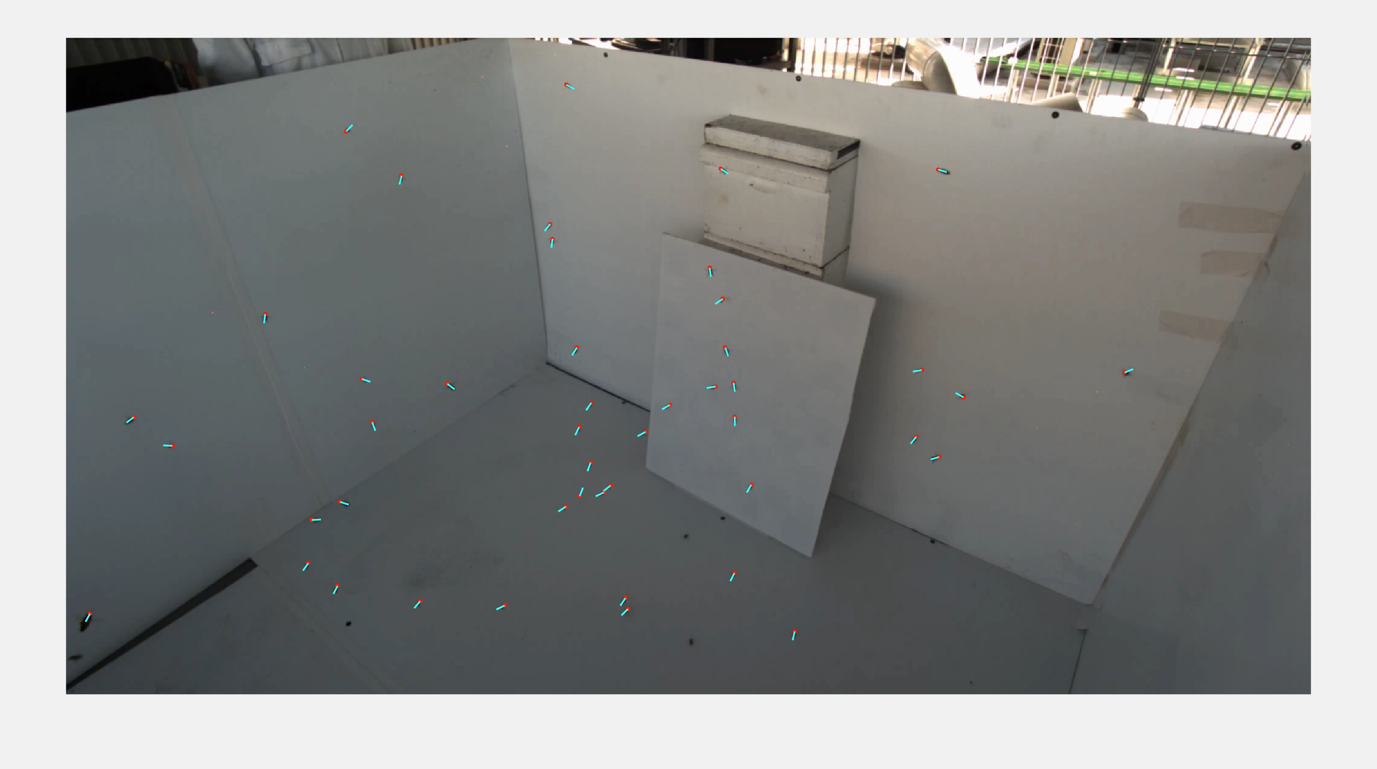

Movie S1- S2. shows the superimposed 2D trajectories of the bees in each camera view. Videos S1 and S2 are from dataset #1. The head position of each bee is indicated by a circular dot in red. The body orientation of each bee is illustrated by a thin line attached to each bee’s head position in cyan. Each bee’s trajectory is shown in different color. For better visualization, the line showing the path traced by a bee in its previous 5 frames are only shown in the video. **Please double click on the video thumbnail to play the video.**

*RECONSTRUCTION OF THREE-DIMENSIONAL TRAJECTORIES OF HONEYBEES FLYING IN HIGH-DENSITY AERIAL ENVIRONMENTS*

Mandiyam Y. Mahadeeswara^1^ and Mandyam V. Srinivasan^1,2^

*SUPPLEMENTARY FILE 2*

*THREE DIMENSIONAL TRAJECTORIES OF HONEYBEES IN THE RECOREDED BEE CLOUD DATA*

*The movie S3 and S4 show the three-dimensional trajectories of all bees flying in the two digitised bee cloud data. The head position and body orientation of each bee in frame is not shown to preserve the clarity of the plot. However, the head position and body orientation of each bee in its last frame of its flight is shown as red dot and black line in the plot, respectively.* **Please double click on the video thumbnail to play the video.**

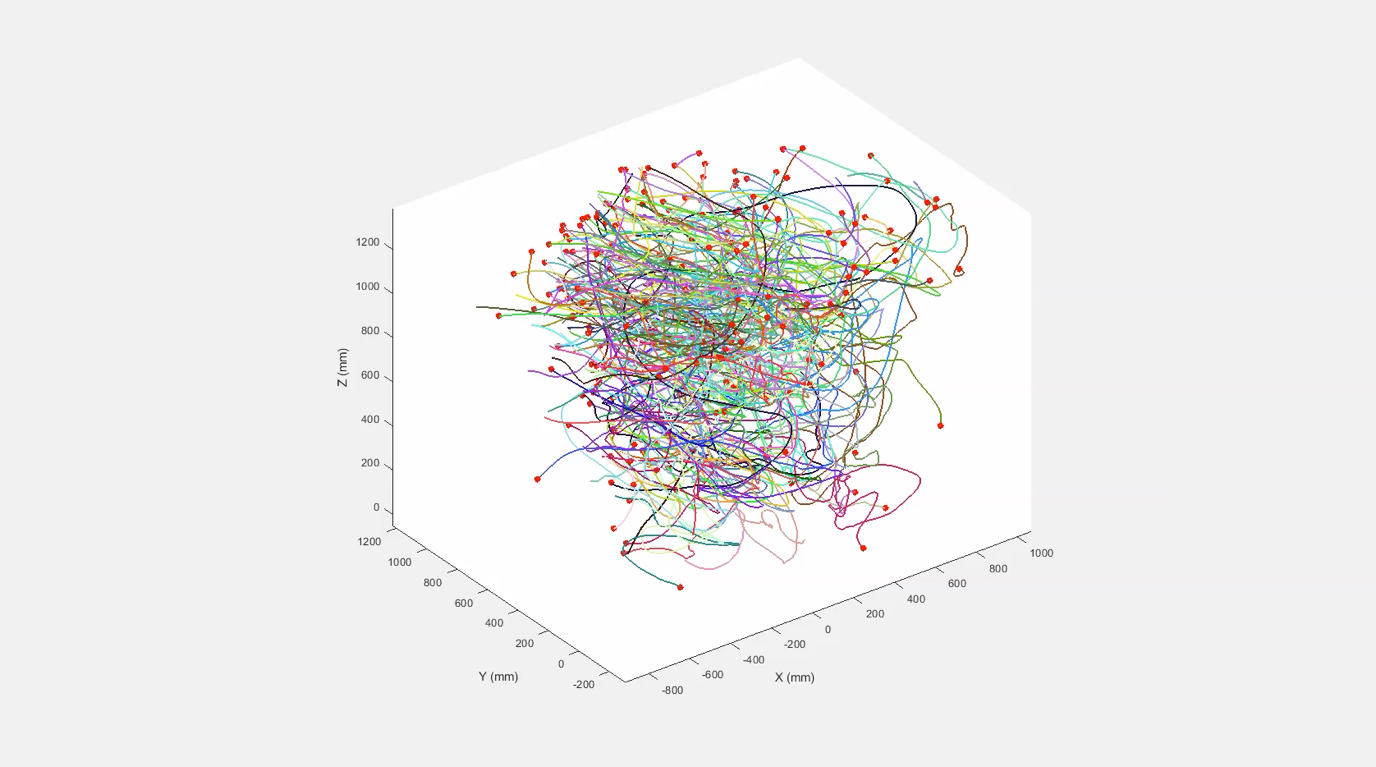
(S3)

(S4)

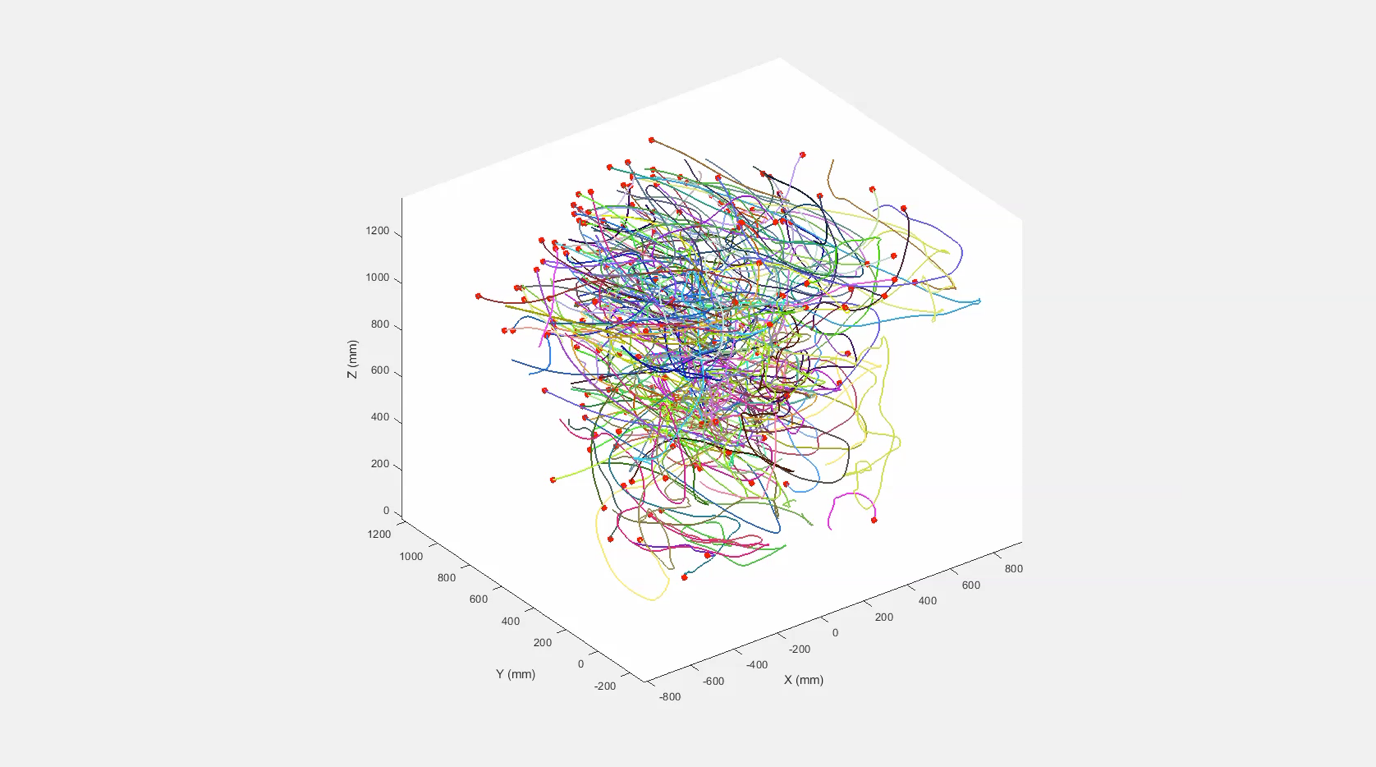

Movie S3-S4: The final three-dimensional trajectories of all bees from Dataset 2. Each color represent individual trajectory of a bee. The head position and body orientation of each bee in its last frame of its flight is shown as red dot and black line in the plot, respectively. **Please double click on the video thumbnail to play the video.**

*RECONSTRUCTION OF THREE-DIMENSIONAL TRAJECTORIES OF HONEYBEES FLYING IN HIGH-DENSITY AERIAL ENVIRONMENTS*

Mandiyam Y. Mahadeeswara^1^ and Mandyam V. Srinivasan^1,2^

*SUPPLEMENTARY FILE 3*

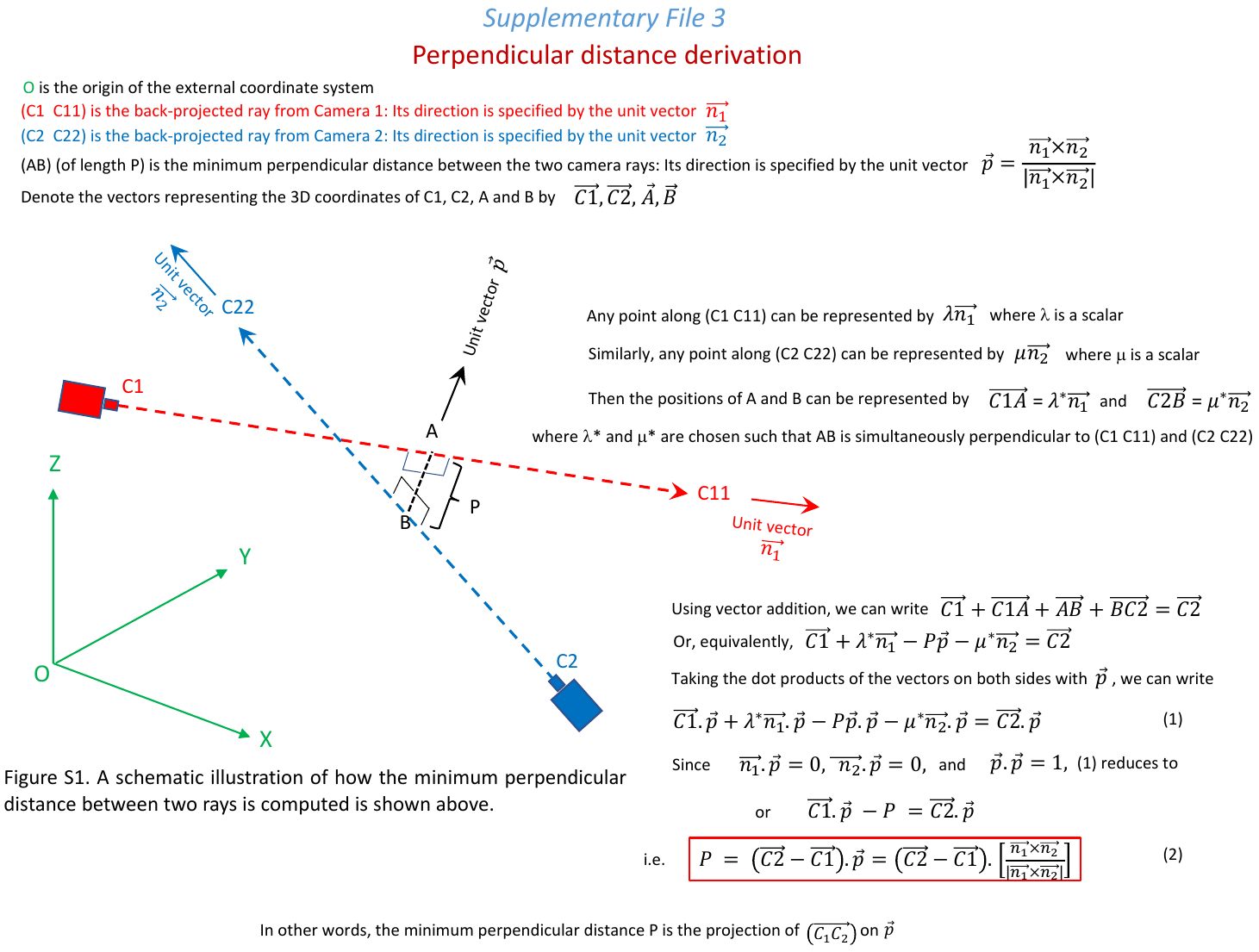
